# Guidance for the design and analysis of cell-type specific epigenetic epidemiology studies

**DOI:** 10.1101/2024.11.06.621949

**Authors:** Emma M Walker, Emma L Dempster, Alice Franklin, Anthony Klokkaris, Barry Chioza, Jonathan P Davies, Georgina ET Blake, Joe Burrage, Stefania Policicchio, Rosemary A Bamford, Leonard C Schalkwyk, Jonathan Mill, Eilis Hannon

**Affiliations:** Department of Clinical and Biomedical Sciences, University of Exeter Medical School, University of Exeter Medical School, University of Exeter, Barrack Road, Exeter, Devon, EX2 5DW, UK; Italian Institute of Technology, Center for Human Technologies (CHT), Genova, Italy; School of Life Sciences, University of Essex, Wivenhoe Park, Colchester, Essex, CO4 3SQ, UK

**Author notes:** **Corresponding author: Eilis Hannon, Department of Clinical and Biomedical Sciences, University of Exeter Medical School, RILD building, Royal Devon & Exeter Hospital, Barrack Road, Exeter. EX2 5DW. UK.**.

**Keywords:** Epigenetic Epidemiology, DNA methylation, complex disease, cell-type specific

## Abstract

Recent studies on the role of epigenetics in disease have focused on DNA methylation profiled in bulk tissues limiting the detection of the cell-type affected by disease related changes. Advances in isolating homogeneous populations of cells now make it possible to identify DNA methylation differences associated with disease in specific cell-types. Critically, these datasets will require a bespoke analytical framework that can characterise whether the difference affects multiple or is specific to a particular cell-type. We take advantage of a large set of DNA methylation profiles (n = 751) obtained from five different purified cell populations isolated from human prefrontal cortex samples and evaluate the effects on study design, data preprocessing and statistical analysis for cell-specific studies, particularly for scenarios where multiple cell types are included. We describe novel quality control metrics that confirm successful isolation of purified cell populations, which when included in standard preprocessing pipelines provide confidence in the dataset. Our power calculations show substantial gains in detecting differentially methylated positions for some purified cell populations compared to bulk tissue analyses, countering concerns regarding the feasibility of generating large enough sample sizes for informative epidemiological studies. In a simulation study, we evaluated different regression models finding that this choice impacts on the robustness of the results. These findings informed our proposed two-stage framework for association analyses. Overall, our results provide guidance for cell-specific EWAS, establishing standards for study design and analysis, while showcasing the potential of cell-specific DNA methylation analyses to reveal links between epigenetic dysregulation and disease.

## Introduction

Recent years have seen increased attention on the role of genetics and gene regulation in development and complex disease (1, 2) facilitated by sequencing and array-based technologies. This includes studies of the epigenome, which encompasses a diverse number of chemical modifications to DNA and nucleosomal histone proteins that directly influence gene expression. In contrast to the genome, the epigenome is highly dynamic, varying across development, between cell-types and in response to the environment. Consequently, this means that careful consideration of study design is required when investigating relationships between epigenetic variation and complex traits (3, 4). The most studied epigenetic modification in the context of human health and disease is DNA methylation (5, 6), which involves the addition of a methyl group to a cytosine. Most existing association studies leverage the high-throughput nature of microarrays to profile DNA methylation (DNAm) at hundreds of thousands of positions across the genome, meaning it is feasible both practically and financially to identify disease-associated variation for large sample numbers.

It is well established that a DNAm profile is primarily defined by tissue or cell type (7-11). Therefore, the choice of tissue for profiling influences both the analytical results obtained and the nature of conclusions that can be drawn from these. Profiling the primary affected tissue, e.g. blood for immune-related traits and the prefrontal cortex for neuropsychiatric and neurodegenerative diseases, is pertinent for making mechanistic inferences about how DNAm variation contributes to a particular trait. However, even these ‘bulk’ tissues represent a heterogeneous mix of cell types. Each of these cell types has its own DNAm profile, with the resulting profile of the bulk tissue being an aggregate of those from the constituent cell types. As the proportion of each cell type within a sample can vary across individuals, systematic differences in cellular proportions that correlate with the phenotype of interest may manifest as differences in the overall DNAm profile (12). To minimise potential false positive associations, quantitative covariates that capture the cellular composition of each sample are typically included in statistical analyses (12). The major caveat of analysing DNAm in bulk tissue is that it does not enable the identification of which cell types are affected by the detected differences. In addition, subtle changes or differences in rarer cell types may be missed as they compete against the background signal from more abundant cell types. Elucidating which cell type(s) are affected by DNAm differences is critical for determining the genes and biological processes associated with specific complex traits and ultimately identifying novel targets for preventing and treating disease.

The case for generating cell-specific DNAm profiles to facilitate cell-specific analyses of epigenetic variation in disease is compelling. Although methods for single-cell DNA methylation profiling have been developed (13-15), these approaches are not currently amenable to large-scale analyses of human disease. Instead, techniques like fluorescence activated nuclei sorting (FANS) and laser capture microdissection can be used to isolate purified cell populations from bulk tissue prior to genome-wide assays and have been applied to tissues such as whole blood (9, 16) and cortex (17-19). Many datasets generated from these methods are small, aimed primarily at generating reference profiles for characterisation of those cell-types or as input for reference-based deconvolution algorithms that estimate the cellular composition from bulk tissue profiles (20). There are, however, cases where these data have been used for epigenetic epidemiology to identify variation in DNAm associated with disease (19, 21-25).

Although the primary tissue for a particular disease may be obvious, the specific cell type involved is often less clear with cell-specific epigenome-wide association studies (EWAS) typically including multiple isolated cell types. In these scenarios, the objective differs from the traditional bulk tissue EWAS. Not only is the goal to identify loci in the genome where differences in DNAm correlate with the outcome of interest, but additionally to characterise whether the difference manifests across multiple cell types or is specific to a particular cell-type. This will require a change in analytical approach because existing statistical approaches based on standard linear regression are potentially inadequate for analysing these data. First, unlike most traditional bulk tissue analyses, multiple samples will be profiled for each individual. This threatens a major assumption of linear regression, that the observations are independent. Second, the statistical framework must capture differences between cases and controls that could be present in all cell types or are only present in an individual (or a subset of) cell types, by estimating case-control differences per cell type and assess whether these are statistically consistent.

In this manuscript, we evaluate the effects on study design, data preprocessing and statistical analysis for cell-specific studies of DNAm, particularly where multiple cell types are considered, by taking advantage of a large DNAm dataset including five different purified cell populations isolated from prefrontal cortex. First, we describe necessary extensions to established quality control pipelines that ensure that the isolation of purified cell populations has been successful. Second, we investigate the effect on statistical power of an association study that uses cell-specific DNAm data. Third, we assess the impact of how the data are normalised on the variance in DNAm. Finally, we evaluate multiple statistical frameworks for analysing cell-specific DNAm data, using two simulation scenarios: a null association study and an association study with differentially methylation positions introduced. With these results we provide guidance for the field in anticipation of future cell-specific EWAS with the objective of establishing standards for how these studies are designed, analysed and interpreted.

## Methods

### Isolation of neural nuclei from post-mortem brain tissue

Post-mortem tissue from 287 adult donors (aged 18-108 years old) was provided from multiple brain banks from the UK, Canada and US (Cambridge, Edinburgh, Stanley, King’s College London, Harvard, UCLA, Oxford, Miami, Douglas Bell, Pittsburgh and Mount Sinai Brain Banks). Tissue was collected under approved ethical regulation at each centre and transferred to our care through Materials Transfer Agreements. Post-mortem human prefrontal cortex (PFC) samples were processed using a FANS protocol developed by our group (26). Details of the gating strategies we implemented are shown in **Supplementary Figure 1**.

### Methylomic profiling

DNA was extracted from frozen nuclei aliquots using a modified proteinase K-based extraction method developed by our group specifically for these sample types (27). 500 ng of genomic DNA from each sample was treated with sodium bisulfite using the Zymo EZ-96 DNA Methylation-Gold™ Kit (Cambridge Bioscience, UK) according to the manufacturer’s standard protocol. All samples were then processed using the EPIC 850K array (Illumina Inc, CA, USA) according to the manufacturer’s instructions, with minor amendments and quantified using an Illumina iScan System (Illumina, CA, USA). Individuals were randomised and sorted fractions from the same individual and FANs gating run were processed on the same BeadChip, where within a BeadChip the location of each fraction was randomised. Altogether, 293 Total (unsorted), 290 NeuNPos (NeuN+), 286 SOX10Pos (NeuN-/SOX10+), 27 IRF8Pos (NeuN-/SOX10-/IRF8+), 274 DoubleNeg (NeuN-/SOX10-) and 15 TripleNeg (NeuN-/SOX10-/IRF8-) nuclei samples were processed.

### DNAm data preprocessing and quality control

DNAm data was loaded into R (version 3.6.3) from idat files using the package bigmelon (28). These data were processed through a bespoke quality control pipeline developed for cell-specific DNAm data. The pipeline is structured into three stages:

Stage 1 - confirming the quality of the DNAm data

Stage 2 - confirming the correct individual

Stage 3 - confirming the correctly labelled cell type

Full details on the steps within these stages can be found in **Supplementary Text S1**. After stringent quality control 751 samples were retained: 218 Total, 164 DoubleNeg, 182 NeuNPos, 168 Sox10Pos, 12 IRF8Pos and 7 TripleNeg samples.

### Comparison of normalisation strategies

For this analysis we limited the comparison to only include cell types that had more than 100 samples (NeuNPos, Sox10Pos and DoubleNeg). The data from the three cell-type fractions were normalised using the *dasen*() function in the wateRmelon R package (29) in two different ways. First, all samples from all cell types were normalised as a single dataset. Second, normalisation was performed for each cell type separately. We compared different normalisation strategies using three quantitative metrics proposed in the original wateRmelon manuscript (29) where, low scores indicate a higher signal-to-noise ratio and a more effective normalisation strategy. Details of these metrics can be found in **Supplementary Text S2**. In addition, we quantified the magnitude of transformation post normalisation using the *qual*() function from the bigmelon R package(28) for each sample.

### Power calculations

To quantify the effect on statistical power of isolating purified populations of nuclei, we performed a series of power calculations using the function *pwr.t.test()* from the R package pwr (30). We consider the scenario with a binary outcome (i.e. case control study), using a two-sample t-test to compare the means of the two groups. We profiled the effect of varying sample size or mean difference between groups on statistical power for each cell type separately. For this, we calculated the cell-specific SDs for each site. The significance level was held constant and set to our previously calculated experiment-wide threshold of P< 9×10^-8^(31).

To get a power estimate for the overall study of 846,232 autosomal sites for a specific scenario, we calculated the cumulative percentage of sites that had that level of statistical power. To reduce the computational burden of performing > 800,000 power calculations per cell type, we implemented a binning method. This takes advantage of the fact that many sites have similar SDs, and therefore the power calculations do not need to be calculated for each site individually. Sites were assigned to 500 bins of equal size (i.e. same number of sites per bin) with each bin containing sites that have similar SDs. The mean of the SDs across sites within each bin is used to calculate Cohen’s d which is input into the power calculation and the resulting power statistic is applied to all sites in that bin. When analysing how the mean difference affected statistical power, we fixed the sample size at either 100 or 200 per group. We then evaluated a range of increasing mean differences (0.001, 0.002, 0.003, …, 0.1). When analysing how sample size affected statistical power, we fixed the mean difference between groups at either 0.02 or 0.05. We then evaluated 100 equally spaced sample sizes ranging from 0 up to the sample size needed to detect a minimum of 85% of sites with at least 80%, rounded to the nearest 100. Note that in these results we report the mean difference for DNA methylation measured as percentage points (i.e. bounded between 0 and 100) rather than as a proportion (i.e. bounded between 0 and 1).

### Simulation study to assess statistical frameworks for cell-specific EWAS

To assess the different analytical frameworks, we implemented two simulation scenarios. First, we generated 100 null association studies by randomly assigning samples to be either cases or controls. Second, starting with each simulated null association study above, we introduced a fixed number of differentially methylated positions at a random subset of sites, predetermined to be either common (i.e. affect all cell types) or specific to a single cell type. Further details on the simulation scenarios can be found in **Supplementary Text S3**.

For each simulation we compared four regression frameworks:

i. Linear regression model for each cell type separately (‘ctLR’)
ii. Linear regression model using all samples from all cell types (‘allLR’).
iii. Mixed effects linear regression model using all samples from all cell types (‘MER’).
iv. Clustered robust regression model using all samples from all cell types (‘CRR’).

All models included age, sex and brain bank as covariates. Further details can be found in **Supplementary Text S4**. For each simulation and regression model, we recorded the number of significant associations at epigenome-wide significance (9×10^-8^) (31) as well as three discovery thresholds (1×10^-7^, 1×10^-6^, 1×10^-5^), classifying these as either a false positives or true positives.

## Results

### Novel quality control pipeline for processing cell-specific DNA methylation data

Quality control of raw DNAm data prior to statistical analysis is critical to minimise technical variation. While several analysis pipelines already exist (29, 32-34), most of these were not developed for studies with multiple measures (i.e. from different cell types) per individual. We have developed a custom pipeline for cell-sorted DNAm data, structured into three stages (**Figure 1**; see **Methods**). Briefly, stage 1 confirms that the DNAm assay has worked and generated high quality data. Stage 2 confirms that the sample matches their labelled individual by checking concordance with sex and genotype information. These first two stages are common to most preprocessing pipelines whereas the final stage of the pipeline is tailored to the cell-sorted dimension of the dataset to confirm that FANS isolation was successful. Leveraging the fact that cell-type identify is the primary source of variation in DNAm profiles (8, 10, 35), the major principal components (PC) should cluster the samples by cell type and can therefore be used to predict cell type. Of note, “unsorted samples” where either no antibody was applied, or where the antibody staining failed, will sit in the middle of the scatterplot, reflecting a heterogeneous set of cells. For each cell type, we calculated the mean and SD of the first two PCs to define its average location in the PC space. Using these values we calculated sample-level scores to assess how closely each sample matched the expected cell type profile. Two checks were implemented based on these scores. First, an “isolation efficiency score”, the median score across samples for that individual, was defined to verify that we successfully isolated distinct fractions of nuclei. Visual inspection of the minimum and maximum isolation efficiency score determined a threshold of 5 was appropriate to identify individuals to exclude (**Supplementary Figure 2**). Second, we confirmed that each sample clustered correctly with its labelled cell type, defined as being within two SDs of its cell type’s mean for the first two PCs.

**Figure 1.**
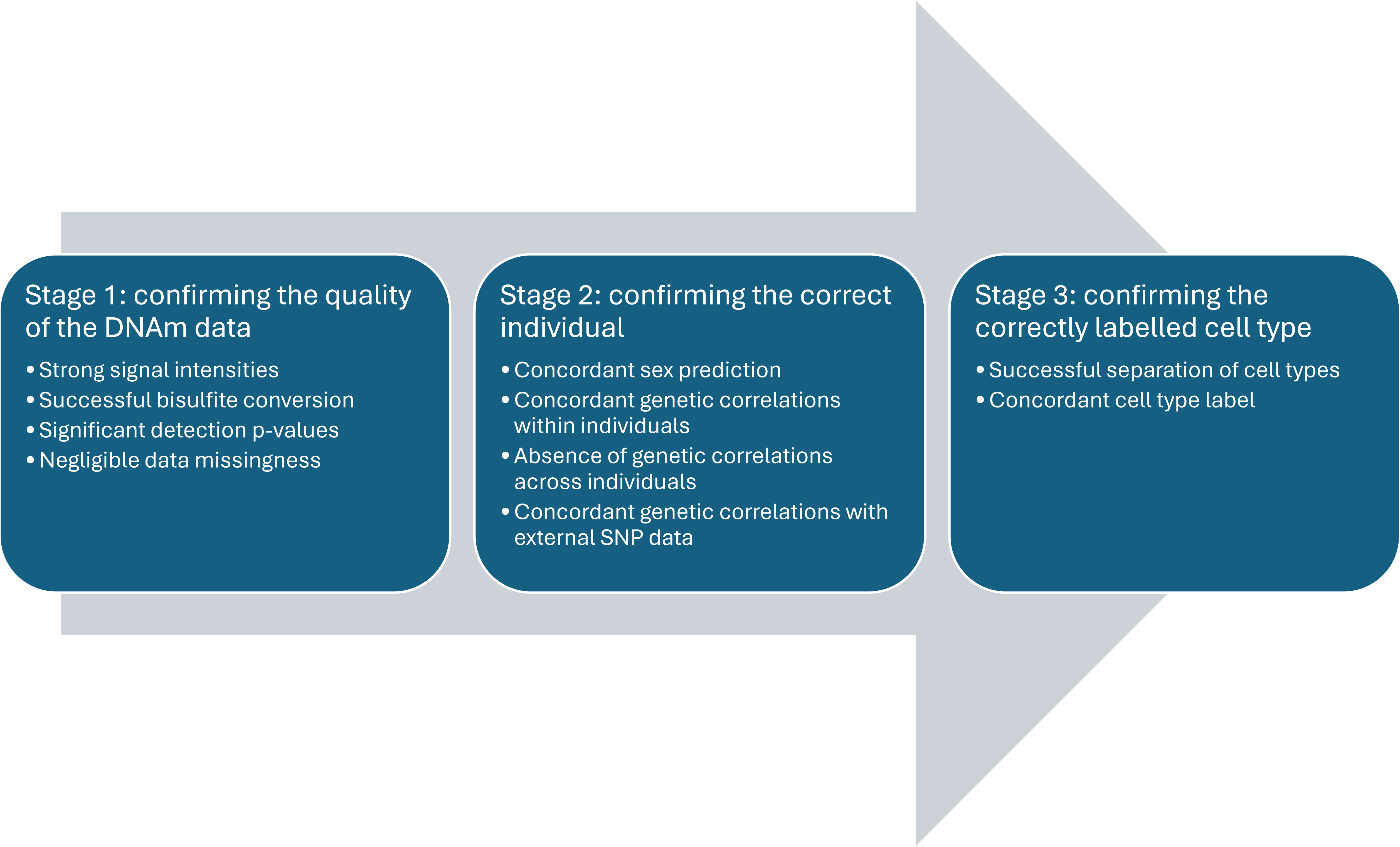
Overview of stages in quality control pipeline for cell-specific epigenetic data.

### Increased sensitivity for detecting differentially methylated positions following normalisation within cell types

After quality control, technical variation from processing samples on different plates needs to be neutralised to minimise confounding and maximise sensitivity. Normalisation achieves this by transforming samples to a common standard, making them quantitatively comparable. While studies report little impact of normalisation strategy on DMP detection in EWAS of single sample types (36, 37), dramatic differences in DNAm profiles between cell types may lead to unpredictable behaviour if these data are forced to become more similar to each other. We compared the effect of normalising across all samples from all cell types together to normalising separately within each cell type using an established framework based on three metrics that can be used to rank the performance of each strategy (29). Across all three metrics, normalising the data separately for each cell type was associated with better performance (**Table 1**). Furthermore, quantifying the effect of normalisation (**Supplementary Figure 3**), we observe that for NeuNPos samples when the samples are normalised with the other cell types, a much larger transformation is made on average to each site (mean = 0.039, SD = 0.016), compared to when they are normalised only with NeuNPos samples (mean = 0.027; SD = 0.011). DoubleNeg and Sox10Pos samples were associated with similar levels of transformation when normalised separately (DoubleNeg mean = 0.033; SD = 0.013; Sox10Pos mean = 0.027; SD = 0.012) compared to when normalised altogether (DoubleNeg mean = 0.032; SD = 0.014; Sox10Pos mean = 0.026; SD = 0.011). This is not surprising given we have previously observed that the primary axis of variation in samples sorted from the PFC is captures differences between NeuN positive and NeuN negative samples (35) meaning these samples require the largest manipulation to make them comparable to the rest.

**Table 1.** Summary metrics to compare normalisation strategies for cell-sorted DNAm data. Each column presents the results of a different metric proposed by Pidsley et al to quantitative compare different normalisation algorithms. For each metric a lower value indicates a superior performing normalisation approach.

| Normalisation strategy | Metric |  |  |
| --- | --- | --- | --- |
|  | DMRSE | GCOSE | Seabird |
|  | imprinted regions | SNP probes | X-chromosome inactivation |
| raw unnormalised data | 1.578E-03 | 5.724E-05 | 0.053 |
| normalise all samples from all cell types together | 1.600E-03 | 5.716E-05 | 0.037 |
| normalise each cell type separately | 1.243E-03 | 5.107E-05 | 0.036 |

### Isolating homogeneous populations of cells reduces the number of samples required to reliably detect differentially methylated positions

It is well established that large sample sizes are required to robustly detect epigenetic differences associated with complex disease. We have previously shown that for a small difference in DNAm (2% points between groups), 500 cases and 500 controls is required for 85% of sites to have at least 80% power to detect a true DMP (31). Therefore, it is often assumed that cell sorting is impractical for large scale epigenetic studies but extrapolating from power calculations derived from heterogeneous bulk tissues can be misleading. Much of the variation in bulk tissues arises from differences in cell composition and using purified cell populations removes this variance likely increasing power to detect cell-type-specific differences. We tested this hypothesis by quantifying the probe level variation by cell type to perform power calculations. Comparing the distribution of SDs across autosomal sites by cell type (**Supplementary Figure 4**), lower variation across samples was observed as expected for the NeuNPos (mean = 0.036; SD = 0.023) and Sox10Pos (mean = 0.038; SD = 0.024) fractions relative to the Total samples (mean = 0.046; SD = 0.026). Total contains nuclei from all fractions and therefore can be considered a proxy for bulk PFC tissue. Of note the DoubleNeg (mean = 0.057; SD = 0.033) and IRF8Pos (mean = 0.055; SD = 0.036) fractions exhibit on average higher levels of variation with the TripleNeg (mean = 0.044; SD = 0.029) fraction showing similar levels of variation to Total.

Taking these data forward for power calculations, we can see that this has a positive effect on the sample size requirements (**Figure 2**). For example, to detect at mean difference of 5% points between groups with power of 80% at 80% of probes on the EPIC array, you need 84 samples per group for NeuNPos or 88 samples per group for Sox10+ (**Supplementary Table 1**). In contrast, for the Total samples you need 132 per group, 128 TripleNeg, 204 IRF8Pos or 212 DoubleNeg per group. Instead to detect a 2% difference you need 476 samples per group for NeuNPos compared to 799 samples per group for Total or 1224 samples per group for IRF8Pos. Alternatively keeping the sample size constant at 100 samples per group, we would have 80% power at 80% of sites to detect a mean difference of 4.5% for NeuNPos, 4.7% for Sox10Pos, 5.7% for TripleNeg, 5.8% for Total, 7.3% for IRF8Pos and 7.5% for DoubleNeg between groups. This decreases to a mean difference of 3.1% for NeuNPos, 3.3% for Sox10Pos, 4.0% for TripleNeg, 4.1% for Total, 5.1% for IRF8Pos and 5.2% for DoubleNeg between groups if the sample size is increased to 200 samples per group.

**Figure 2.**
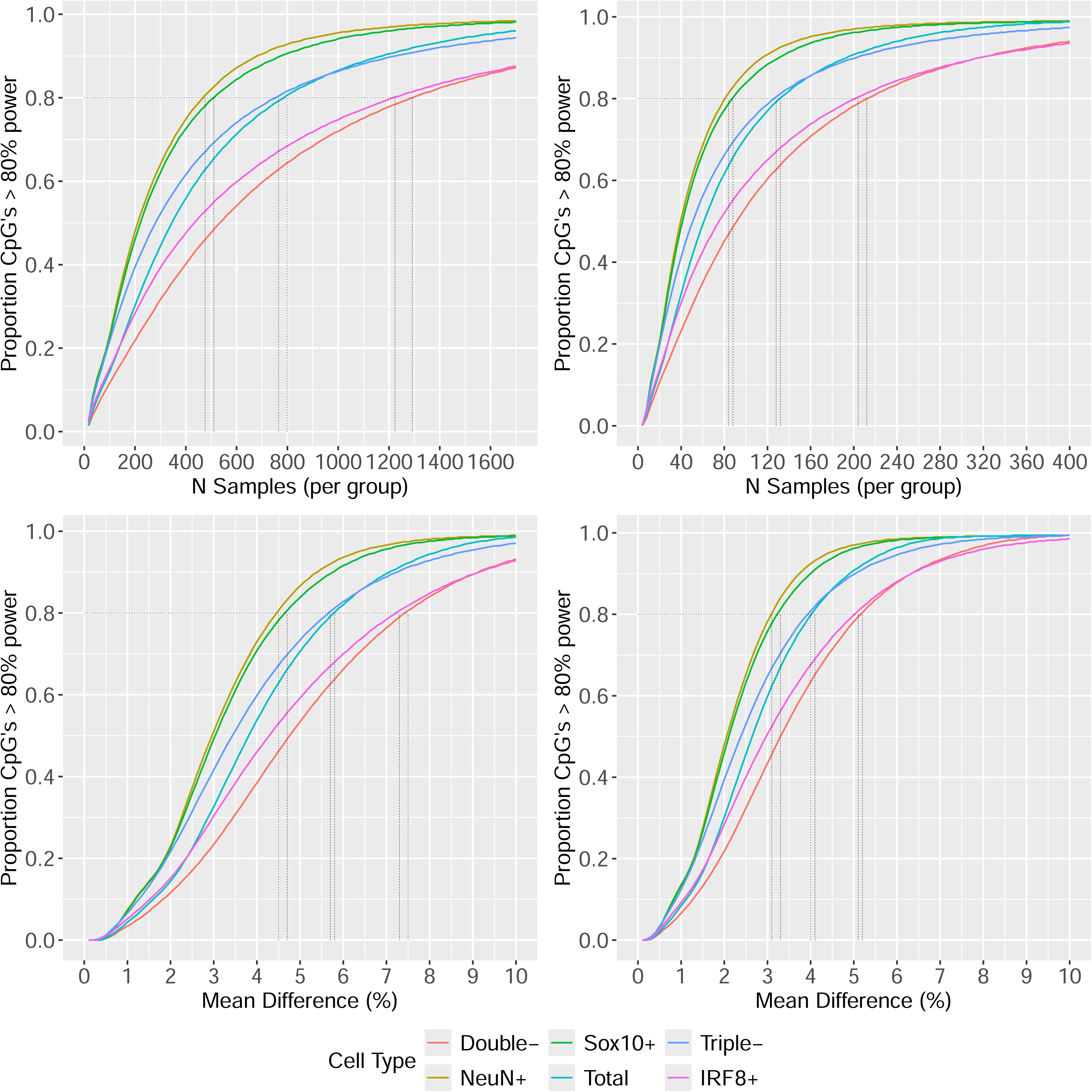
Cell-specific epigenetic association analyses increase power over bulk tissue studies. Presented are power curves for different purified brain cell types, and bulk brain tissue for different experimental parameters. Panels **A** & **B** show line graphs of power (y-axis) against sample size (i.e. the number of samples per group) for mean differences between groups of 5% (**A**) or 2% (**B**). Panels **C** & **D** show line graphs of power against mean difference (%) for fixed sample sizes of 100 cases vs 100 controls (**C**) and 200 cases vs 200 controls (**D**). Each line represents a different brain cell type or bulk brain tissue. Highlighted are experimental parameters under which a minimum power of 80% to detect a difference is observed.

### Cell-specific EWAS are prone to false positives if within individual study design is not adequately controlled for

Given the differences in study design and objective compared to single tissue EWAS, a novel analytical framework is accounts for multiple samples per individual and that allows for differentially methylated positions (DMPs) to affect a subset of cell types. Ordinary least squares linear regression will likely be biased, so a more complex regression model is required, but there are multiple alternatives available. To evaluate the impact of regression model on statistical sensitivity, we designed a simulation framework to compare four different approaches. The first approach (‘ctLR’) is linear regression within each cell type, where there is no correlation between samples to control for. It is a computationally cheap analysis per DNAm site but involves fitting one model per cell type and is unable to determine the cell-specificity of the effect. The second approach (‘allLR’) uses all the data available from all cell types in a linear regression model. This method violates the assumption of independent observations but by aggregating the data together should increase power to detect DMPs that affect multiple cell-types. The third approach (‘MER’) uses a mixed effects regression model to control for the structure within the data. The final approach (‘CRR’) uses a clustered robust regression model that is an alternative approach to account for related samples. The allLR, MER and CRR methods all include both a main effect and interaction terms to capture DMPs impacting different cell types.

When deciding upon an efficient method there are two desirable properties: minimising the false positive rate and maximising the true positive rate. To quantify the false positive rate for each approach we simulated 100 null EWAS (see **Methods**). For regression models that include all data from all samples (allLR, MER & CRR), the significance of the main effect and/or interaction term depends on which cell type a DMP affects, so both terms must be considered when assessing these methods. We observed that the distribution of p-values for both the allLR and MER models is wider with longer tails indicative of smaller mean p-values (**Supplementary Figure 5**). Applying the standard threshold for epigenome-wide significance 9×10^-8^ (31) we can see that the ctLR and CRR are well calibrated with all models identifying a mean of < 0.1 false positives (**Figure 3**). In contrast both the allLR and MER models had increased rates. These were dramatically inflated for the main effect term (allLR mean = 42 (SD = 136), MER mean = 53 (SD = 185) and subtly higher also for the interaction term (allLR mean = 1 (SD = 2); MER mean = 2 (SD = 3)). This highlights that the choice of analysis method is important and can have dramatic effects on the robustness of the results. One mechanism to counteract this, is to derive a significance threshold calibrated appropriately for each method. Calculating the mean 5% family-wise error rate for each method across the 100 simulations (**Supplementary Figure 6**) we found that the significance thresholds for the ctLR and CRR models were comparable to the standard threshold (range: P = 9.7×10^-8^ – 2.8×10^-7^), whereas the thresholds for the allLR and MER models were 2-7 orders of magnitude smaller (range: P = 9.7×10^-16^ – 5.3 × 10^-9^).

**Figure 3.**
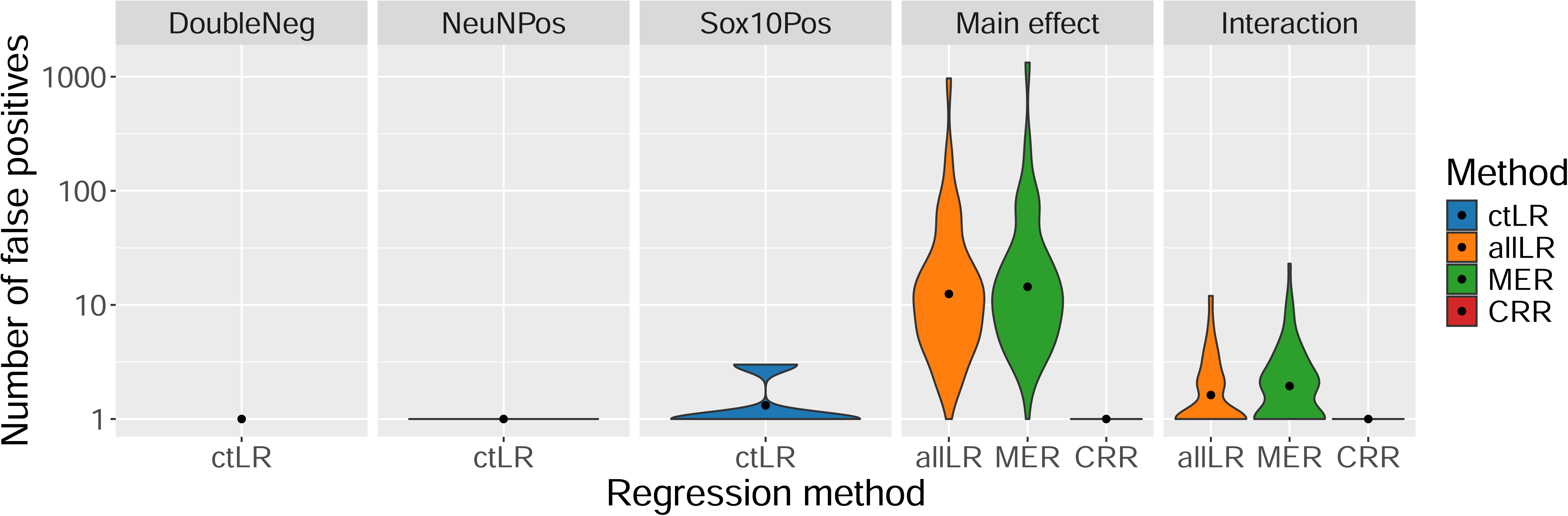
Choice of regression method for cell-specific association analyses is critical to minimise false positive associations. Violin plots of the number of false positive differentially methylated positions (y-axis, log scale) detected (P < 9×10^-8^) for different regression methods from 100 simulated null EWAS. Each violin represents a different statistical analysis and the colour of the violin indicates the regression method. ctLR - within cell-type linear regression; allLR – linear regression with samples from all cell-types; MER – mixed effects regression; CRR – clustered robust regression.

To assess the ability of each method to detect true positive associations we also implemented a second simulation scenario where a fixed number of DMPs were introduced with effects that were either common to all three cell types or specific to just one. Considering DMPs with a mean difference of 5% between cases and controls, the ctLR in NeuNPos samples had the highest true positive rate (mean = 0.76, SD = 0.35; **Figure 4**), slightly higher than the MER (mean = 0.74, SD = 0.09), ctLR in Sox10Pos samples (mean = 0.73, SD = 0.34) and allLR (mean = 0.72, SD = 0.09). There was then a drop in performance to the CRR method (mean = 0.66, SD = 0.10) followed by another drop in performance to the ctLR in DoubleNeg samples (mean = 0.49, SD = 0.26). The disparity in performance with the ctLR method for different cell types fits with the results of the power calculations and highlights how the increased variation of DoubleNeg samples can impact the ability to detect true associations. Generally, performance did not depend on whether a DMP was common to all cell types or specific to just one, although it is notable that the CRR model had a higher true positive rate for detecting cell-specific DMPs (mean = 0.68, SD = 0.13) than either the MER (mean = 0.65, SD = 0.14) or allLR (mean = 0.60, SD = 0.15). All methods performed slightly worse than the ctLR in either NeuNPos (mean = 0.72, SD = 0.36) or Sox10Pos (mean = 0.68, SD = 0.38) samples. This pattern of results was maintained when the mean difference between cases and controls was reduced to 2%, albeit with reduced true positive rates in general (**Supplementary Figure 7**; range of mean true positive rates for all DMPs = 0.12-0.27). Of note, there was no meaningful effect of the number of DMPs on the performance of the regression method (**Supplementary Figure 8**). The proportion of cell-type specific DMPs didn’t affect the true positive rate of the ctLR, but interestingly the rate did decrease as the proportion increased for the allLR and MER methods but increases for the CRR method.

**Figure 4.**
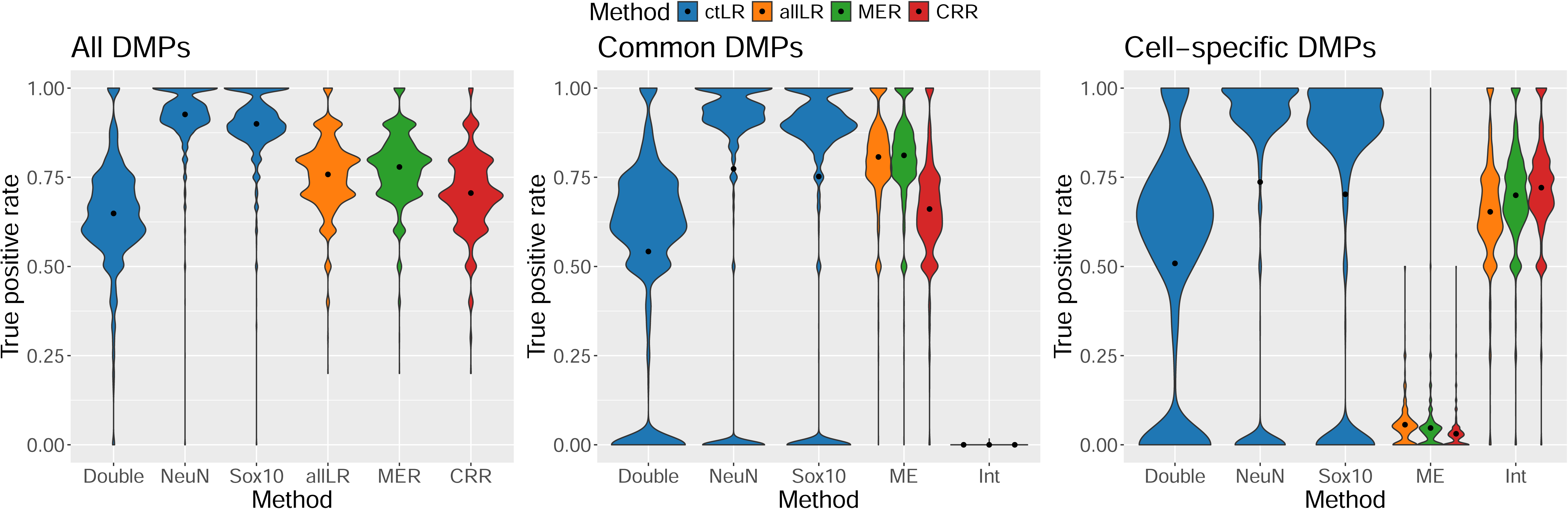
Violin plots comparing true positive rates across cell-specific regression methods. Violin plots of true positive rate (P < 9×10^-8^) calculated for different regression methods across 1800 simulated EWAS with 10, 100 or 1000 differentially methylated positions introduced with mean difference of 5% between groups. A proportion of differentially methylated positions (0,0.2,…,1) were allocated to be specific to one cell type (“cell-specific”), and the rest affected all cell types (“common”). Panels represent true positive rates calculated for **A**) all differentially methylated positions, **B**) cell-specific differentially methylated positions only and **C**) common differentially methylated positions only. DMPs - differentially methylated positions; ctLR - within cell-type linear regression; allLR – linear regression with samples from all cell-types; MER – mixed effects regression; CRR – clustered robust regression.

Many EWAS fail to identify DMPs at the standard epigenome-wide significance threshold attributing this to a lack of statistical power. It is not uncommon to see “discovery thresholds” used, which are slightly relaxed significance levels, to uncover some potential associations that may in future larger studies, transpire to be true positives. To date, no one has assessed how valuable this approach is or whether it is dominated by false positives. First, considering the number of false positives for a range of discovery thresholds (1×10^-7^, 1×10^-6^, 1x-10^-5^) for the methods that had minimal false positives at epigenome-wide significance, ctLR and CRR, the mean false positive count remained less than 1 up to a threshold of < 10^-6^ and increased to 4-10 at p-value threshold of < 10^-5^ (**Supplementary Table 2**). Meanwhile, the methods that already had elevated mean false positive counts continued to accumulate increasing numbers of false positives from 43 to 656 for allLR and 55 to 785 for MER at p-value < 10^-5^. Secondly, considering the true positive rate, this increases as the significance threshold is relaxed for all methods, albeit to different degrees (**Supplementary Table 3**). There are smaller gains for the methods that were already associated with high rates, e.g. ctLR in NeuNPos samples increases from 0.91 (SD = 0.08) at 9×10^-8^ to 0.95 (SD = 0.06) at 10^-5^ with bigger gains for methods that had lower rates e.g. CRR increases from 0.66 (SD = 0.10) at 9×10^-8^ to 0.78 (SD = 0.09) at 10^-5^.

## Discussion

We present the first quantitative assessment of study design, quality control and statistical analysis for cell-specific DNAm association studies. Using data from 5 purified populations of nuclei obtained by FANS from post-mortem cortical tissue we use both empirical power calculations and simulations to determine a statistically robust approach to not only identify positions in the genome associated with an outcome, but to characterise which cell types these differences affect. Determining the cell types affected by differences in DNAm is the next critical step in enhancing our understanding of the role of epigenetics in health and disease.

Our motivation parallels challenges in single-cell transcriptomics, which similarly involves multiple replicates per sample (on a substantially greater scale) and identifying the cell-specificity of differential gene expression. Analyses of empirical and synthetic datasets in that domain showed that many proposed methodologies, including as we did mixed effects models, failed to account adequately for the variability between replicates and consequently were associated with inflated rates of false positives (38). Concerns regarding the proliferation of unsound analyses contributing to misguided subsequent scientific studies have prompted significant methodological research and ongoing discourse in the field. As cell-specific EWAS are in their infancy, we propose that now is the critical time to establish a framework that promotes statistically robust analyses facilitating the generation of meaningful and accurate biological insights.

Balancing the need to minimise the false positive rate and maximise the true positive rate we propose the following analytical strategy for EWAS of multiple cell types. First, we recommend performing a within cell type association analysis using linear regression. From each regression model (one per cell type), significant DMPs can be identified with low risk of false positive associations, subject to appropriate study design and adjustment for confounders. The caveat with this approach is that it cannot determine whether the identified DMP is cell type-specific or affects multiple cell types. Cross-referencing DMP lists across cell types may reveal overlaps, but a lack of overlap does not confirm absence of effect, as it could result from sampling variation or limited statistical power. Therefore, a second stage of the analysis is required to confirm whether the effect is common to all cell types by testing for differences or heterogeneity across cell types. We propose that a mixed effects model is used in this scenario with an interaction term and random effect to control for multiple samples per individual. While this method was associated with a high number of false positives, we are not proposing using it to discover DMPs but simply to characterise the cell-type specificity of the effect. We additionally investigated the impact of using a “discovery threshold” but this offered minimal benefit in identifying additional true positives. If required, we recommend not exceeding 10^-6^, as this kept the number of false positives below 1.

One of our key take home messages is that isolating populations of cells may not need to be performed in as many samples as would be needed for a bulk tissue EWAS. Our power calculations found that for some purified populations, there were substantial gains in statistical power for detecting DMPs. Consequently, in studies with limited samples (e.g. post-mortem brain tissue), cell-sorting could be a valuable strategy to boost statistical power and maximise the impact of a limited resource. Within specific cell types it is anticipated that effect sizes will be greater, enhancing power further. However, for some cell-sorted fractions, such as IRF8Pos fractions (microglia enriched) and TripleNeg fractions (astrocyte enriched) there was an increase in variation and therefore a decrease in power. This suggests that within these fractions, further isolation may be required to not only increase the probability of finding genuine associations, but also to clarify which cell types show increased variation in DNA methylation.

The findings we present should be considered in light of the following limitations. First, we only consider a case-control study design, however, the use of a regression framework, means our approach can be adapted for more complex study designs. There is no obvious reason that our results would not also hold for continuous phenotypes for example. Second, while our data only consists of neural purified cell populations, we believe the overarching conclusions of our study are generalisable to cell types from other tissues. For those interested in DNAm EWAS of brain cell types, we have developed an R package, CellPower, for the community to perform power calculations for brain cell-specific EWAS. Third, we only considered a limited number of parameter combinations for DMPs and sample size. We believe this is sufficient to address the questions we posed, but caution should be applied when extrapolating from the specific true positive and false positive rates we report to other EWAS scenarios. Fourth, we used a microarray to quantify DNAm with limited coverage of CpGs. There are other sequencing based technologies that can be used to profile DNAm more extensively across the genome that generate estimates with different statistical properties (39) and it may be that our results do not extend to these other technologies.

In conclusion, our analyses provide a valuable insight into the potential of cell-specific DNA methylation association analyses and provide a benchmark against which future studies can be evaluated in terms of their study design, quality control pipelines and statistical analysis.

## Data Availability

Raw and processed DNA methylation data are available from GEO under accession number GSE279509.

## Code Availability

All code for the results presented in this manuscript are available at https://github.com/ejh243/BrainFANS. The bespoke quality control pipeline we describe can be found at https://github.com/ejh243/BrainFANS/tree/master/array/DNAm/preprocessing, codes for the analyses can be found at https://github.com/ejh243/BrainFANS/tree/master/array/DNAm/analysis/methodsDevelopment. We have also made our data and method for cell-specific EWAS power calculations available as a standalone R package which is available at https://github.com/ew367/CellPower/tree/main.

## Ethics approval and consent to participate

Ethical approval for the study was granted by the College of Medicine & Health Research Ethics Committee under application reference number 6524714.

## Acknowledgements

We acknowledge the supply of samples from; The Cambridge Brain Bank, covered by current REC approval NRES 10/HO308/56; the Quebec Suicide Brain Bank at Douglas Mental Health University Institute, Canada; the MRC funded University of Edinburgh Brain & Tissue Bank; the Harvard Brain Tissue Resource Centre, which is supported by HHSN-271-2013-00030C; Brain Endowment Bank, at Miller School of Medicine, University of Miami; The Mount Sinai NBTR (NIH Brain and Tissue Repository), JJ Peters VA Medical Center; the Oxford Brain Bank, supported by the Medical Research Council (MRC), the NIHR Oxford Biomedical Research Centre and the Brains for Dementia Research programme, jointly funded by Alzheimer’s Research UK and Alzheimer’s Society; The Neuropathology Brain Bank at the University of Pittsburgh, School of Medicine Department of Psychiatry; The Stanley Medical Research Institute Brain Collection courtesy of Drs. Michael B. Knable, E. Fuller Torrey, Maree J. Webster, and Robert H. Yolken; The Human Brain and Spinal Fluid Resource Center (HBSFRC) NIH Neurobiobank. This study was supported by the National Institute for Health and Care Research Exeter Biomedical Research Centre. The views expressed are those of the author(s) and not necessarily those of the NIHR or the Department of Health and Social Care. For the purpose of open access, the author has applied a CC BY public copyright licence to any Author Accepted Manuscript version arising from this submission.

## Key Points

- We propose an analytical framework for cell-specific DNA methylation data that incorporates bespoke quality control metrics and not only performs an association analysis to identify differentially methylation positions but that also characterises whether it affects multiple cell types or is specific to a particular cell-type.
- Consideration must be given to the most appropriate analytical model for these data, as even methods that theoretically adjust for correlations between samples, may be associated with inflated rates of false positives.
- Isolating purified cell types can lead to increases in statistical power to identify differentially methylated positions and therefore this might be an effective way of maximise the value of a limited resource such as post-mortem brain tissue samples.

## Biographical Note

Dr Hannon is an Associate Professor in Bioinformatics in the Complex Disease Epigenetics Group at the University of Exeter Medical School and currently holds anEPSRC Research Software Engineering Fellowship.

