## Supplementary Figure for "Guidance for the design and analysis of cell-type specific epigenetic epidemiology studies"

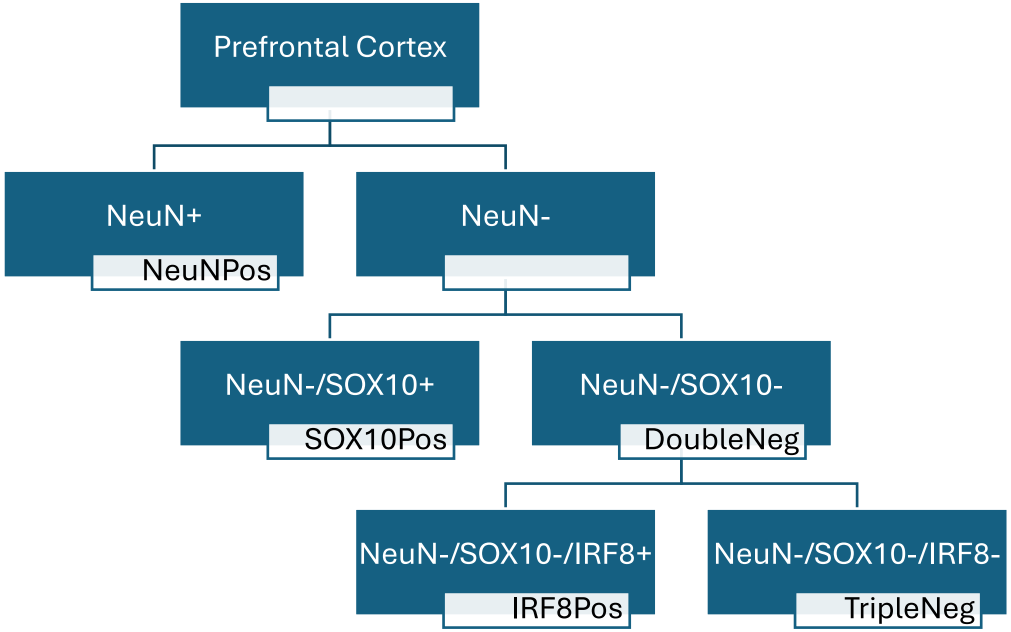


**Supplementary Figure 1**. Schematic of gating strategy to isolate brain cell types. The blue boxes state the combination of antibodies used and the white boxes give the labels used to refer to that sample type in the manuscript.

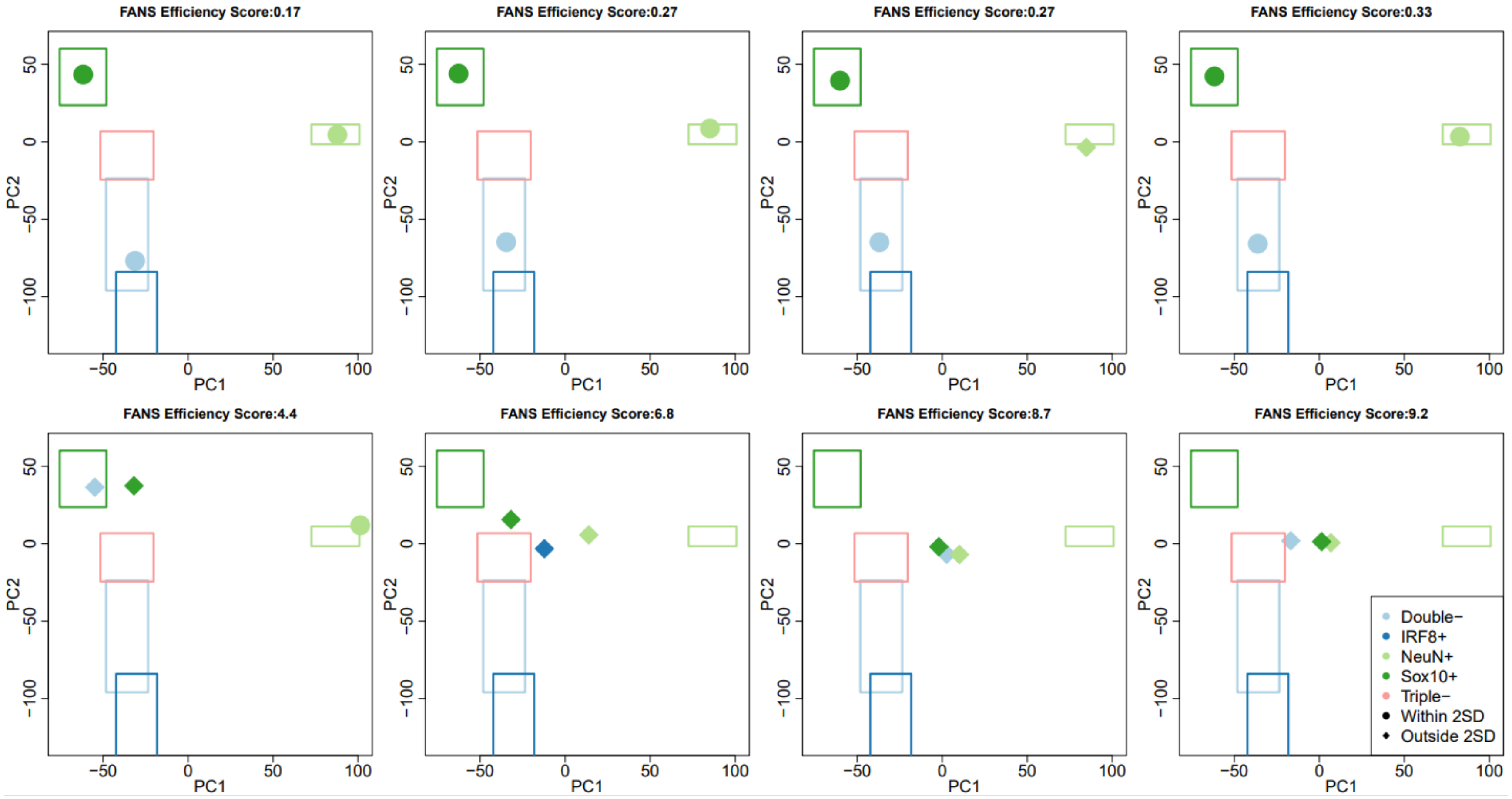


**Supplementary Figure 2.** Validation of the FANs efficiency score for verifying a successful FANs sort. Scatterplots of the first two principal components (PC) of DNA methylation data where each point represents a sample coloured by their labelled cell type. The coloured box represents the location of all samples labelled as that cell type, centred on the mean for that PC and extending two standard deviations in each direction to indicate how similar each sample is to the average profile of that cell type. Each panel includes the samples from a single individual. The top row includes examples of individuals where the FANs sort was successful at obtaining distinct populations of nuclei. The bottom row includes examples of individuals where the FANs sort was less successful at obtaining distinct populations of nuclei.

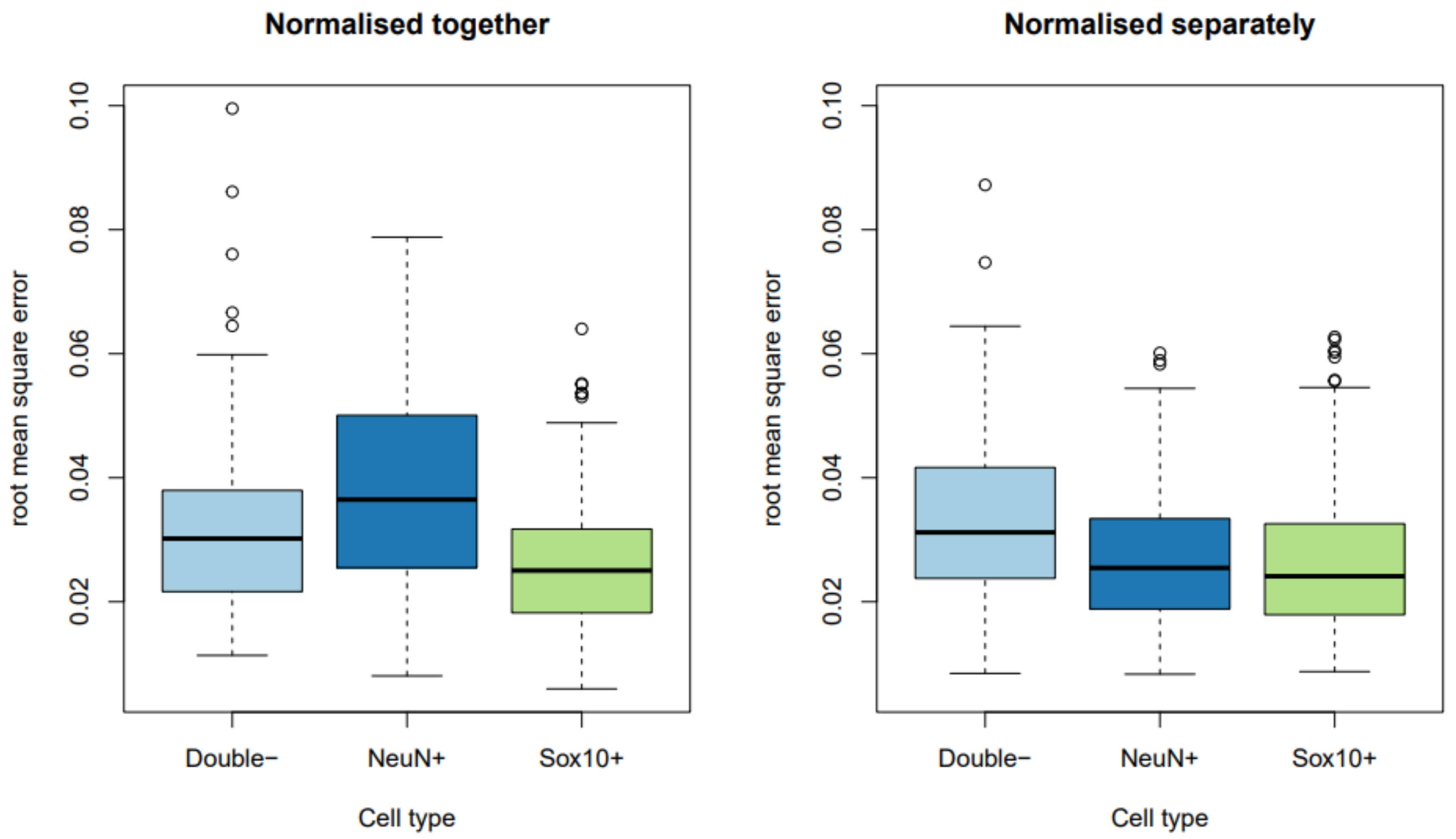


**Supplementary Figure 3** Comparison of the magnitude of the normalisation effect on samples by normalisation strategy. Violin plots of the distribution of normalisation effects across samples, measured as the root mean square error between DNA methylation levels before and after normalisations across all sites. Samples are grouped by cell type. Each panel presents the effect of a different normalisation strategy: left) all samples normalised together; right) samples normalised separately for each cell type.

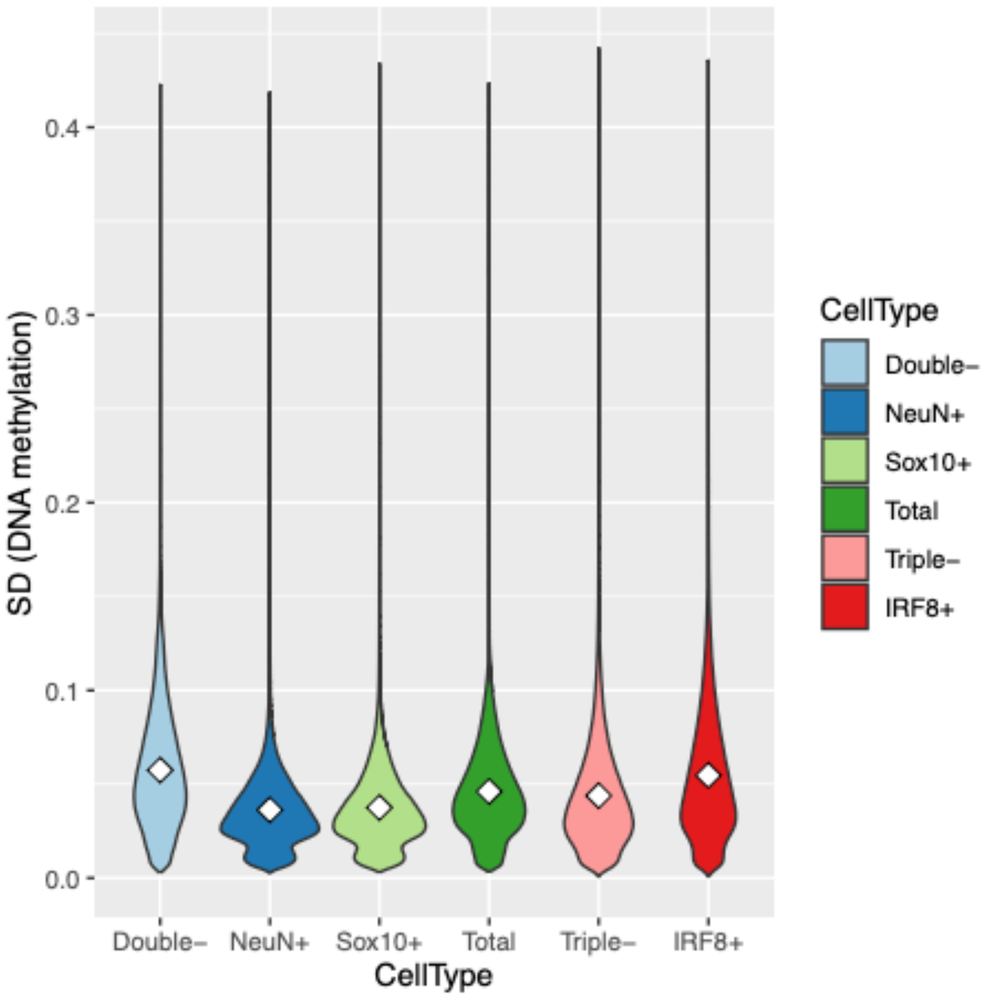


**Supplementary Figure 4.** Violin plots of the distribution of the standard deviation (SD) of DNA methylation across 866,529 autosomal probes on the EPIC v1 array split by cell type.

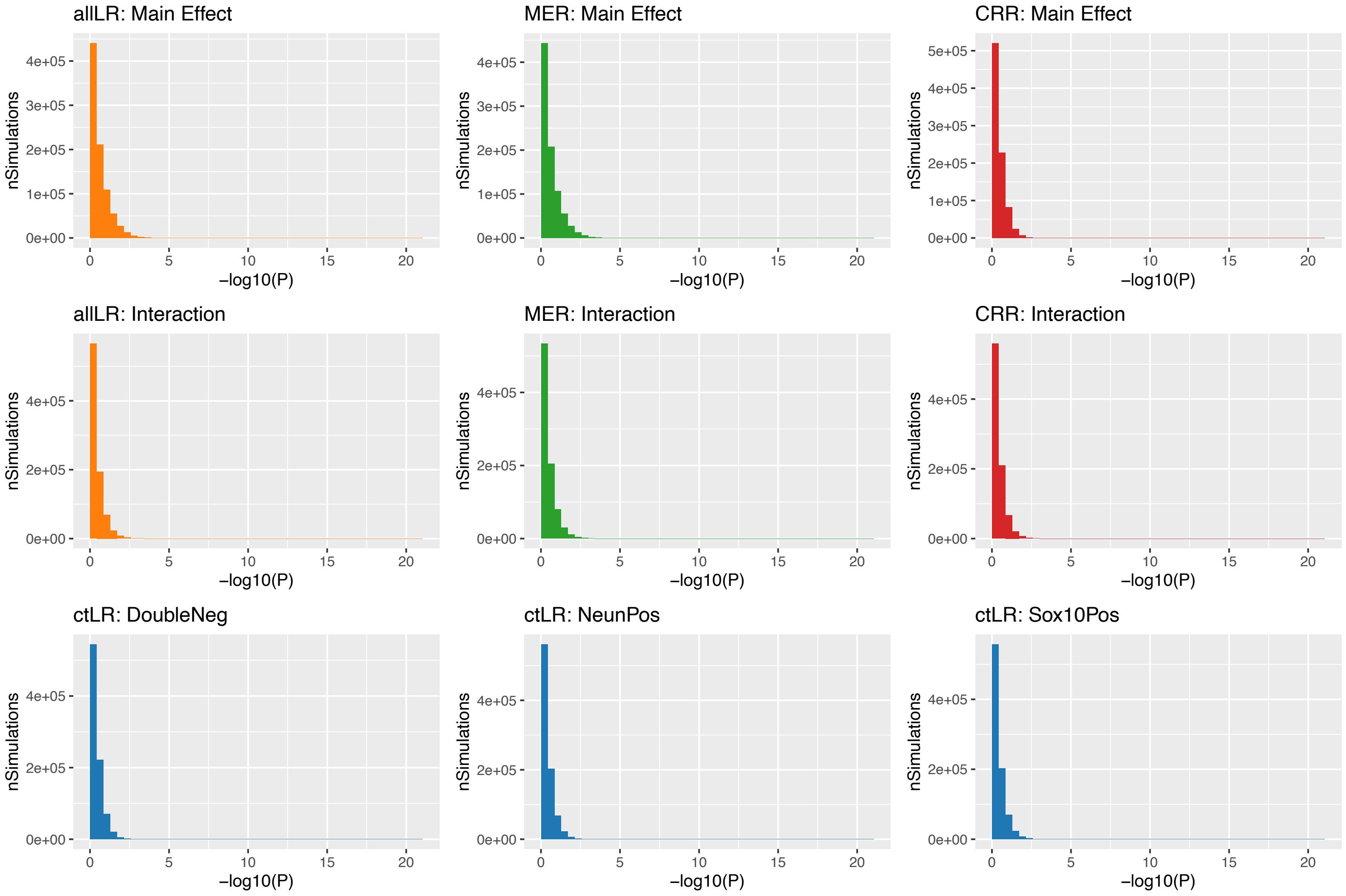


**Supplementary Figure 5.** Histograms of null EWAS p-value distributions. Plotted is the distribution of all -log10(p-values) for all 866529 sites across all 100 simulations, where each panel contains the results from a different method. ctLR - within cell-type linear regression; allLR – linear regression with samples from all cell-types; MER – mixed effects regression; CRR – clustered robust regression.

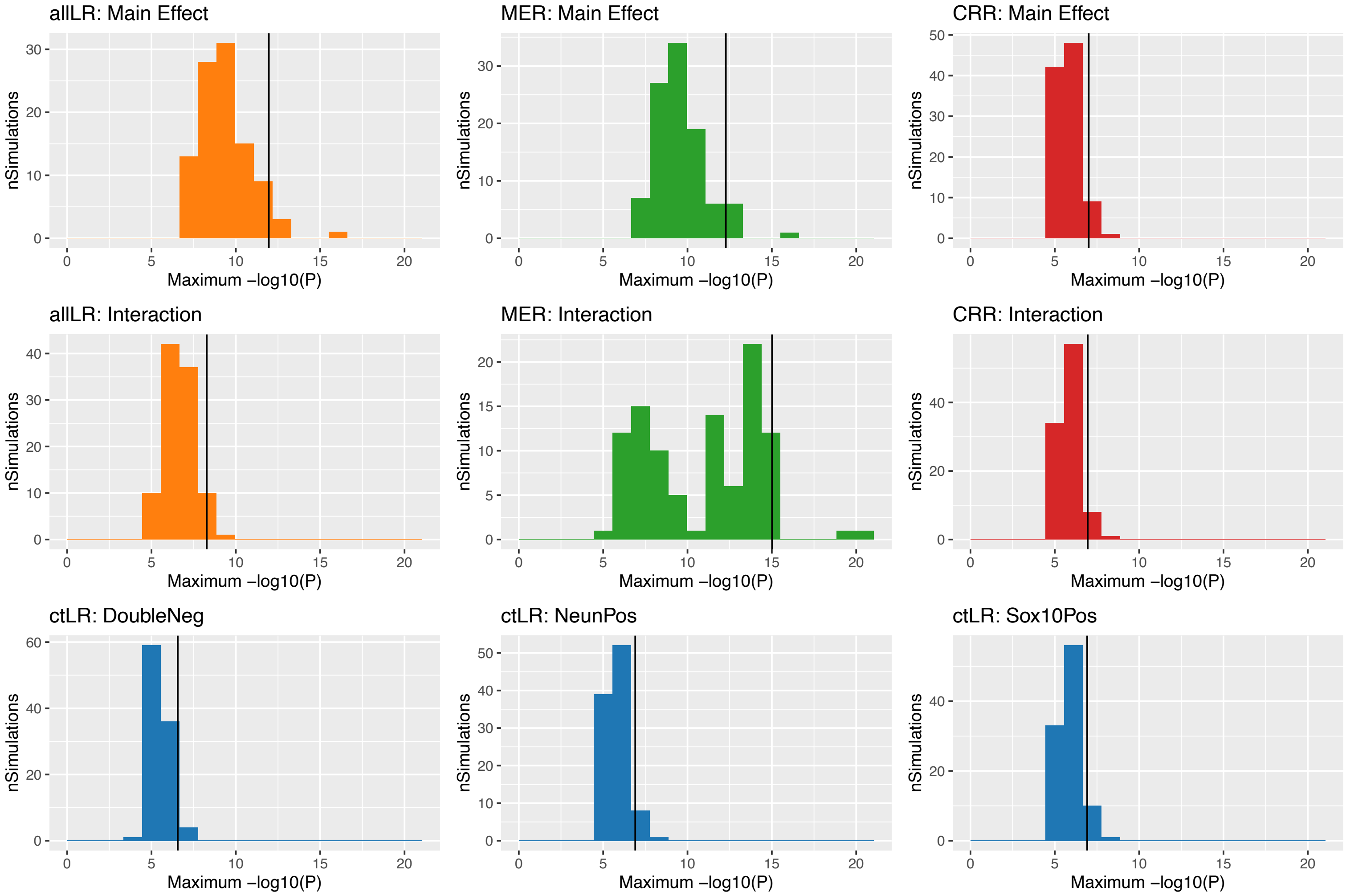


**Supplementary Figure 6.** Histograms of null EWAS minimum p-value distributions. Plotted is the distribution of the 5th percentile -log10(p-values) from 100 simulations calculated across 866,529 sites, where each panel contains the results from a different method. ctLR - within cell-type linear regression; allLR – linear regression with samples from all cell-types; MER – mixed effects regression; CRR – clustered robust regression.

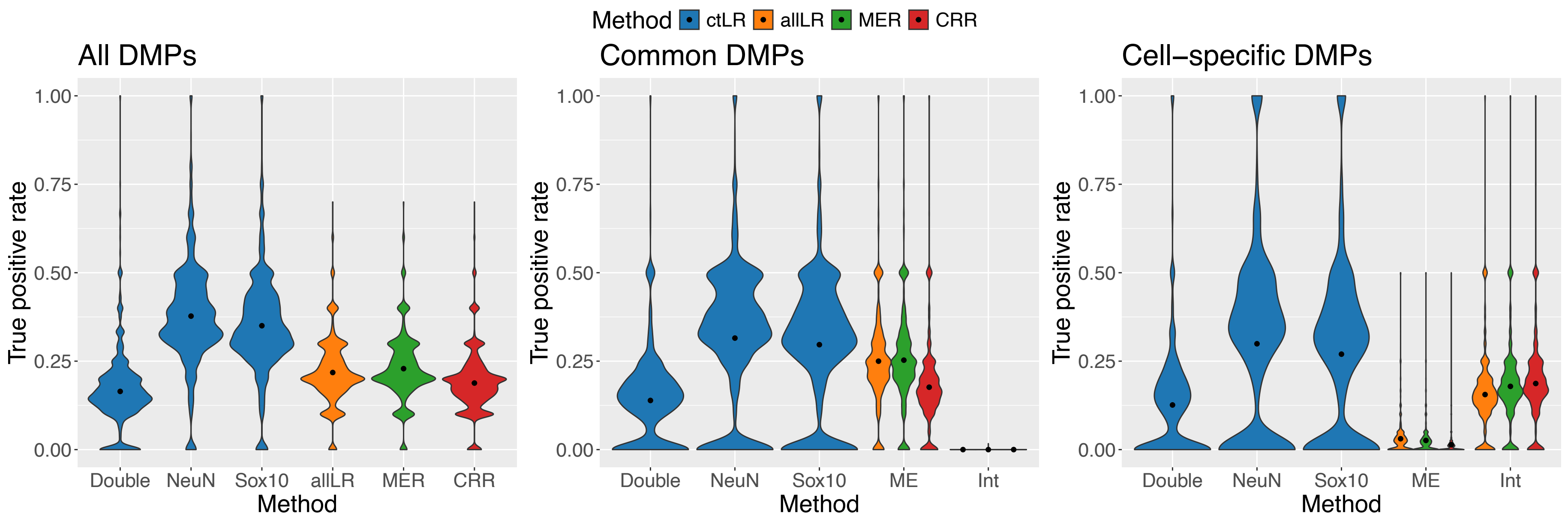


**Supplementary Figure 7. Violin plots comparing true positive rates across cell-specific regression methods.** Violin plots of true positive rate ($P < 9x10^{-8}$) calculated for different regression methods across 1800 simulated EWAS with 10, 100 or 1000 differentially methylated positions introduced with mean difference of 2% between groups. A proportion of differentially methylated positions (0,0.2,…,1) were allocated to be specific to one cell type ("cell-specific"), and the rest affected all cell types ("common"). Each panel presents the true positive rates calculated for all differentially methylated positions, cell-specific differentially methylated positions only and common differentially methylated positions only. DMPs - differentially methylated positions; ctLR - within cell-type linear regression; allLR – linear regression with samples from all cell-types; MER – mixed effects regression; CRR – clustered robust regression.

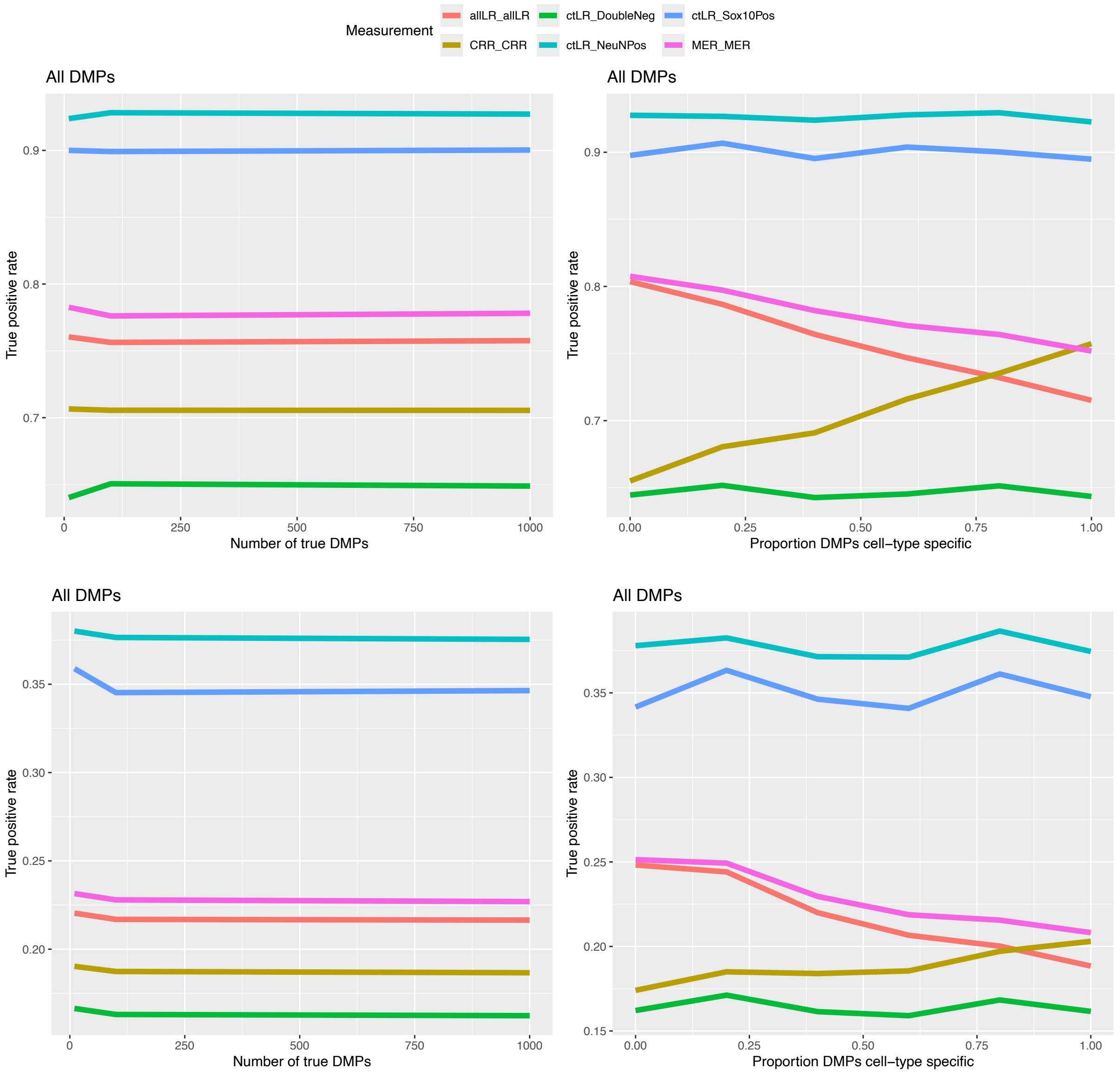


**Supplementary Figure 8**. Line graphs of true positive rate (y-axis) for differentially methylated positions against either number of DMPs (x-axis) or proportion of cell-specific DMPs (**x-axis**). Panels in the top row are taken from simulations where the mean difference between groups was 5% and panels in the bottom row are taken from simulations where the mean difference between groups was 2%. Each line represents a different regression method.
