## Supplementary Text for "Guidance for the design and analysis of cell-type specific epigenetic epidemiology studies"

**Supplementary Methods**

### Supplementary Text S1 - DNAm data preprocessing and quality control

DNAm data was loaded into R (version 3.6.3) from idat files using the package bigmelon (1). These data were processed through a bespoke quality control pipeline developed for cell-specific DNAm data. The pipeline is structured into three stages:

Stage 1: confirming the quality of the DNAm data

i) Checking the methylated and unmethylated signal intensities, excluding samples where this was < 500.
ii) Using the relevant control probes to ensure the sodium bisulfite conversion was successful, excluding any samples with median < 80.
iii) *pfilter* function from wateRmelon package to exclude samples with > 1% of probes with detection P-value > 0.05.
iv) Counting the number of missing values per sample and excluding samples with > 2% probes missing.

Stage 2: confirming the correct individual

v) Prediction of sex from both the X and Y chromosomes by calculating the ratio of the mean intensity from each sex chromosome to the mean intensity on the autosomes. Samples were classified as female if the value from the X chromosome, was > 1.01 or male if the value was < 1 or NA between these values; Samples were classified as female if the value from the Y chromosome was > 0.5 or as male if the value was > 1 or as NA between these values. To confirm the sample identity, the predicted sex had to be consistent across both chromosomes and match the sex recorded in the metadata.
vi) Use of the 59 SNP probes to confirm that samples from the sample individual were genetically identical.
vii) Use of the 59 SNP probes to confirm that samples were not genetically identical to any sample labelled as belonging to another individual.
viii) Comparison of the SNP probes with external matched SNP chip data for that individual, samples were excluded if this correlation was < 0.9.

Stage 3: confirming the correctly labelled cell type

ix) Each sample was compared to the average profile for its labelled cell type to determine how similar it was. Similarity was determined using principal components calculated from the matrix of DNAm levels. For each cell type, the values of the first two principal components were Studentized to identify outliers (which are likely to be due to either mislabelling or suboptimal FANS sorting), excluding those above a threshold of 1.5. The mean and standard deviation (SD) of the first two principal components was then calculated across the remaining samples for each cell type to represent the average profile. These were then used to calculate sample level scores that captured the similarity of the observed sample and the expected profile for the cell type it was labelled as. This was defined as the value of the principal component for that sample minus the cell type mean divided by the cell type SD. The value represents the number of SDs from the mean that sample is, where lower values are desirable. The values from the first two principal components were combined into a single score by taking the maximum, referred to here as the maxSD score. If FANS sorting worked well (i.e. all antibodies were stained and gated accurately), samples should be close to their relevant average profile resulting in a series of low maxSD scores for that individual. Conversely, if sorting failed to isolate the relevant cell types, samples would be heterogeneous mixtures of cells and sit in the middle of the PC space reflected by large maxSD scores. Individual level metrics referred to as the “isolation efficiency score” were calculated as the median across all the maxSD scores for that individual. Using the median highlights unsuccessful sorts rather than instances where one or two samples are affected/mislabelled. All samples for any individual with an isolation efficiency score > 5 were excluded.
x) We recalculated the Studentized values prior to recalculating the cell type means and SD for the first two principal components. Samples that were more than two SDs away from the mean in either of the first two principal components were excluded.

After stringent quality control 751 samples were retained: 218 Total, 164 DoubleNeg, 182 NeuNPos, 168 Sox10Pos, 12 IRF8Pos and 7 TripleNeg samples.

### Supplementary Text S2 – Details of metrics for comparison of normalisation strategies

i) ***DMRSE:*** This metric evaluates sites within imprinted differentially methylated regions (iDMRs) where β values are expected to be 0.5 due to uniparental methylation. The SD across all iDMRs sites (n = 203) is divided by the square root of the number of samples.
ii) ***GCOSE:*** This metric evaluates 58 common SNPs on the array, where β values should follow a trimodal distribution across genotypes. For each SNP, k-means clustering (k = 3) is applied and the sum of squares for each cluster is calculated, summed across SNPs, divided by the sample size, and further divided by the square root of the total number of samples. The mean of the three scores is taken to provide a single statistic.
iii) ***Seabird:*** This metric evaluates X-chromosome sites which have sex-based differences due to X-chromosome inactivation. A t-test p-value, comparing male and female DNAm levels, is used to predict X-chromosome location. A receiver operating characteristic analysis quantifies the prediction accuracy as 1 minus the area under the curve. To improve computational efficiency, a random subset of 200,000 sites (4,173 on the X chromosome) was included.

### Supplementary Text S3 – Details of the simulation study to assess statistical frameworks for cell-specific EWAS

To assess the different analytical frameworks, we implemented two simulation scenarios. For this analysis we limited the comparison to only include cell types that had more than 100 samples (i.e. NeuNPos, Sox10Pos and DoubleNeg).

First, we generated multiple null association studies by randomly assigning samples to be either cases or controls. As there is no reason why the randomly assigned cases should be any more similar to each other than the controls, there should be no genuine differences between the groups. Therefore, any significant differences can be assumed to be false positives. This process was performed 100 times.

Second, starting with the random case control assignment for each simulated null association study above, we introduced a fixed number of differentially methylated positions (DMPs; n = 10, 100, 1000) at a random subset of sites. A proportion (0, 0.2, 0.4,…,1) of these were then randomly selected to be either common (i.e. affect all cell types to the same extent) or specific to a single cell type, where the affected cell type was selected at random. This was achieved by adding the specified mean difference (0.02 or 0.05) the DNAm values for that site to all assigned cases plus a randomly sampled error ~N(0,0.005). This scenario enables us to assess how effective each method is at detecting true positives and whether this differs by the nature of the difference between groups.

### Supplementary Text S4 - Regression frameworks considered in the simulation study

For each simulation we compared four regression frameworks, fitting six different regression models. In all frameworks below, age was included as a continuous variable (estimated from the Cortical Epigenetic Clock (2) as it was not available for all individuals) and sex and brain bank were included as categorical variables.

i) Linear regression model for each cell type in turn (‘ctLR’; total 3 regression models, 1 per cell type). In this model, as there is only one sample per individual per cell type, there is no violation of the assumption that observations are independent. For each cell type the following regression model was fitted for each DNAm site:

$$DNAm \sim case status+age+sex+brain bank$$

ii) Linear regression model using all samples from all cell types (‘allLR’), with covariates to adjust for the mean difference in DNAm between cell types and an interaction term, to capture both common and cell type specific DMPs from a single regression framework. In this model, while there are multiple samples per individual, there is nothing in the model to control for this potential violation of the assumption. The following regression model was fitted for each DNAm site, where i denotes individual and j denotes cell type:

$${DNAm}_{i,j}\sim{case status}_{i}+{cell type}_{j}+{case status}_{i}*{cell type}_{j}+ {age}_{i}+ {sex}_{i}+ {brain bank}_{i}$$

iii) Mixed effects linear regression model using all samples from all cell types (‘MER’), with covariates to adjust for the mean difference in DNAm between cell types and an interaction term, to capture both common and cell-type-specific DMPs from a single regression framework. In this model, a random intercept is included to adjust for multiple samples per individual, and a separate random intercept for brain bank. Case status, cell type, age and sex are included as fixed effects. This model was fitted using the lme4 (3) and lmerTest (4) R packages. The following regression model was fitted for each DNAm site, where i denotes individual and j denotes cell type:

$${DNAm}_{i,j}\sim{case status}_{i}+{cell type}_{j}+{case status}_{i}*{cell type}_{j}+ {age}_{i}+ {sex}_{i}+ {individual}_{i}+ {brain bank}_{i}$$

iv) Clustered robust regression model using all samples from all cell types (‘CRR’), with covariates to adjust for the mean difference in DNAm between cell types and an interaction term, to capture both common and cell type specific DMPs from a single regression framework. In this model each individual is treated as a cluster. The data are pooled across clusters assuming a consistent effect across all individuals and the standard errors of the coefficient estimates (case status, cell type, age, sex and brain bank) are subsequently adjusted using the Huber-White sandwich estimator to account for the potential similarity of samples from the same individual. This model was fitted using the plm (5) and lmtest (6) R packages. The following regression model was fitted for each DNAm site, where i denotes individual and j denotes cell type:

$${DNAm}_{i,j}\sim{case status}_{i}+{cell type}_{j}+{case status}_{i}*{cell type}_{j}+ {age}_{i}+ {sex}_{i}+ {brain bank}_{i}$$
